# Expectations and Motor Signals Shape Auditory Evoked Responses in an Independent Manner

**DOI:** 10.1101/2025.08.07.669075

**Authors:** Batel Buaron, Roy Mukamel

**Affiliations:** Sagol School of Neuroscience and School of Psychological Sciences, Tel-Aviv University

## Abstract

Voluntary actions are usually accompanied by sensory consequences that evoke neural responses in relevant sensory regions. These evoked responses are different when compared to those evoked by otherwise identical physical stimulation not associated with preceding actions (sensory attenuation). A common model suggests that sensory attenuations are caused by efference signals associated with the binding of actions with sensory outcomes. Nevertheless, the information encoded in such signals is debated, with some theories suggesting they convey predictive information while others suggest they convey global motor information regardless of specific outcome expectations. To address this debate, we recorded EEG data while participants (n=30) learned to associate motor or visual cues with corresponding tones. Following 8 repetitions of cue-tone learning, participants successfully learnt the association (>84% correct). At the neural level, the amplitude of auditory evoked responses (N100) decreased across repetitions, as the binding between cues and tones strengthened. In addition, N100 amplitude was attenuated when preceded by motor vs, visual cues. Most importantly, we did not find an interaction effect between repetition and cue type, suggesting a similar sensory attenuation effect across repetitions. Specifically, we show significant sensory attenuation even at the first repetition, when no expectation could be formed. Thus, our results suggest that the sensory attenuation effect does not change while forming experience with action-outcome contingency. These results have implications for contemporary models of sensorimotor control and predictive mechanisms.

## Introduction

In order to learn audio-motor skills, e.g. playing the piano, a student has to be able to couple between each keypress with its specific note. Previous studies have demonstrated that sounds that are the consequences of actions, for example pressing a key on the piano, are perceived differently relative to hearing the same sounds played by someone else (sensory attenuation; (Weiss et al., 2011; Reznik et al., 2015). Not only perception, but also the neural response for the sound is different in such a case. For example, it was found that actively triggering a sound evokes a smaller auditory response relative to passively listening to the same sound (Baess et al., 2009; Timm et al., 2013).

Despite the wealth of evidence indicating a broad and robust influence of voluntary movements on sensory processing, the underlying mechanisms and their functional relevance remain poorly understood. One prevailing model, the forward model (Wolpert and Miall, 1996), proposes that voluntary actions involve an internal mechanism where executed motor commands are accompanied by an ‘efference copy’ sent to sensory regions. It is hypothesized that by modulating the neural state in sensory regions, these signals shape the processing, and eventually the perception of sensory action-outcomes, resulting in the sensory attenuation effect (Horvath, 2015; Reznik and Mukamel, 2019; Kiepe et al., 2021). However, the specific information represented in such efference signals and their developmental course remain largely unknown.

One account for the information encoded in efference signals proposes that they primarily carry predictive information regarding the properties of expected sensory outcome (indicating ‘what is going to happen and when’ (Wolpert and Flanagan, 2001)). According to this account, differences in evoked responses between actively triggered and passively perceived sensory stimuli are derived from the inherent advantage in predicting the content and timing of outcomes of voluntary actions versus external sensory events (Kiepe et al., 2021). Although most studies examined well established motor-sensory couplings (Baess et al., 2009; Lange, 2011; Hughes et al., 2013), some evidence suggests that sensory evoked responses change after learning new coupling. For example, a previous study showed that modulation of responses for self-triggered sounds in mice auditory cortex increase after learning (Schneider et al., 2018). Results from humans also show that introducing a delay between action and consequence results in decreased auditory evoked response for delays that were experienced vs. unexperienced in previous trials (Elijah et al., 2016). Similar to motor predictions, sensory predictions were also found to modulate neural responses of sensory stimuli (relative to unpredictable sensory events; for review see: (Schroger et al., 2015). Although some studies directly compared motor-sensory and sensory-sensory predictions, there is no conclusive answer as to whether they are supported by two distinct mechanisms. A study comparing sensory attenuation effects between motor and non-motor predictors found that both prediction types resulted in similar perceptual sensory attenuation (Dogge et al., 2019). Other findings indicate that some aspects might still be exclusive to motor–sensory predictions. For example, sensory attenuation for self-generated sounds is reduced (but not abolished) when one controls for temporal prediction (Lange, 2011; Hughes et al., 2013).

Another prominent account of the information encoded in efference signals suggests they convey information regarding the motor action and ownership over the generated outcome, giving rise to the feeling of agency that ‘I am the causal agent controlling this outcome’ (Kiepe et al., 2021). In support of this notion, it was found that schizophrenic patients, whose pathology is characterized by misattribution of agency (deficiency in ascribing the causal relationship between action and outcome), do not show motor-induced sensory attenuation (Shergill et al., 2005; Ford et al., 2007). Studies focusing on neurotypical participants examined the sensory attenuation effect when controlling for stimulus characteristics and/or temporal predictability factors (Weiss and Schutz-Bosbach, 2012; Klaffehn et al., 2019; Storch and Zimmermann, 2022). For example, Weiss and Schutz-Bosbach (2012) showed that perceptual sensory attenuation occurs both when the external stimulus was expected or unexpected, suggesting that sensory attenuation is specifically tied to self-generation. Note that the predictive and motor information accounts are not necessarily mutually exclusive and attenuated responses to the sensory consequences of voluntary actions can result from a combination of the two.

In the current study, we examined the relative contribution of predictions to the sensory attenuation effect, by isolating it from other motor related components. To this end we used EEG to examine the magnitude of auditory-evoked responses (N100) during a learning process, as participants accumulate predictive experience with either motor (button-presses) or non-motor (visual) cues and subsequent sounds. Accumulating experience with cue-sound contingency should change the evoked-response under the prediction account, since the coupling between the action and sensory outcome is strengthened with experience. Under the motor information account, sensory attenuation should remain stable during contingency learning since agency is invariant to experience level.

## Methods

### Participants

Thirty-five participants were recruited for the study. Data from 3 participants were discarded due to poor performance on the task (see below), and additional data from 2 participants were discarded due to technical problems in EEG recording, leaving a total of 30 participants for analysis (12 males, mean age 23.13 years, range 18-29 years). All participants were healthy, right-handed (determined by self-report) and had normal hearing and normal or corrected to normal vision. Participants were naïve to the purposes of the study. The study conformed to the guidelines that were approved by the ethical committee at Tel-Aviv University. All participants provided written informed consent to participate in the study and were compensated for their time by either course credits or 40$.

### Experimental Design and Statistical Analysis

#### Procedure

In order to evaluate how auditory evoked responses change as a function of coupling between predictive cues and auditory tones, participants were engaged in a de-novo learning task while their neural activity was recorded using EEG. Each participant underwent learning with two types of predictive cues – button presses (Motor Cues condition) or visual stimuli (Visual Cues condition). Each experimental trial consisted of two phases: a learning phase followed by a test phase evaluating their learning (see Figure 1A). During the learning phase in the Motor Cues condition, participants had to learn the association between three buttons and three different tones (100ms pure tones including 15ms up and 15ms down ramping). Throughout this phase, participants viewed three empty rectangles presented on the center of the screen. Each time one of the rectangles was colored grey, participants had to press the corresponding button on a keyboard. Pressing the button immediately triggered the tone associated with it (see figure 1B). Throughout the trial, each button was pressed 8 times, resulting in 8 repetitions for each button-sound coupling. Participants were instructed to learn the association between each button and the specific tone it generated. On each trial, a novel mapping of buttons to tones was created (de-novo learning). This design allowed us to directly examine the influence of specific expectations on evoked neural activity by comparing the magnitude of the auditory N100 component between different stages of the learning process. In the first repetition of each trial, participants had no knowledge about the mapping between button and sound, while in the last repetition such an association had been formed (and quantified in the test phase). Therefore, our design allowed us to directly examine the influence of expectations on neural activity by comparing the magnitude of the auditory N100 across different stages of the learning process.

**Figure 1:**
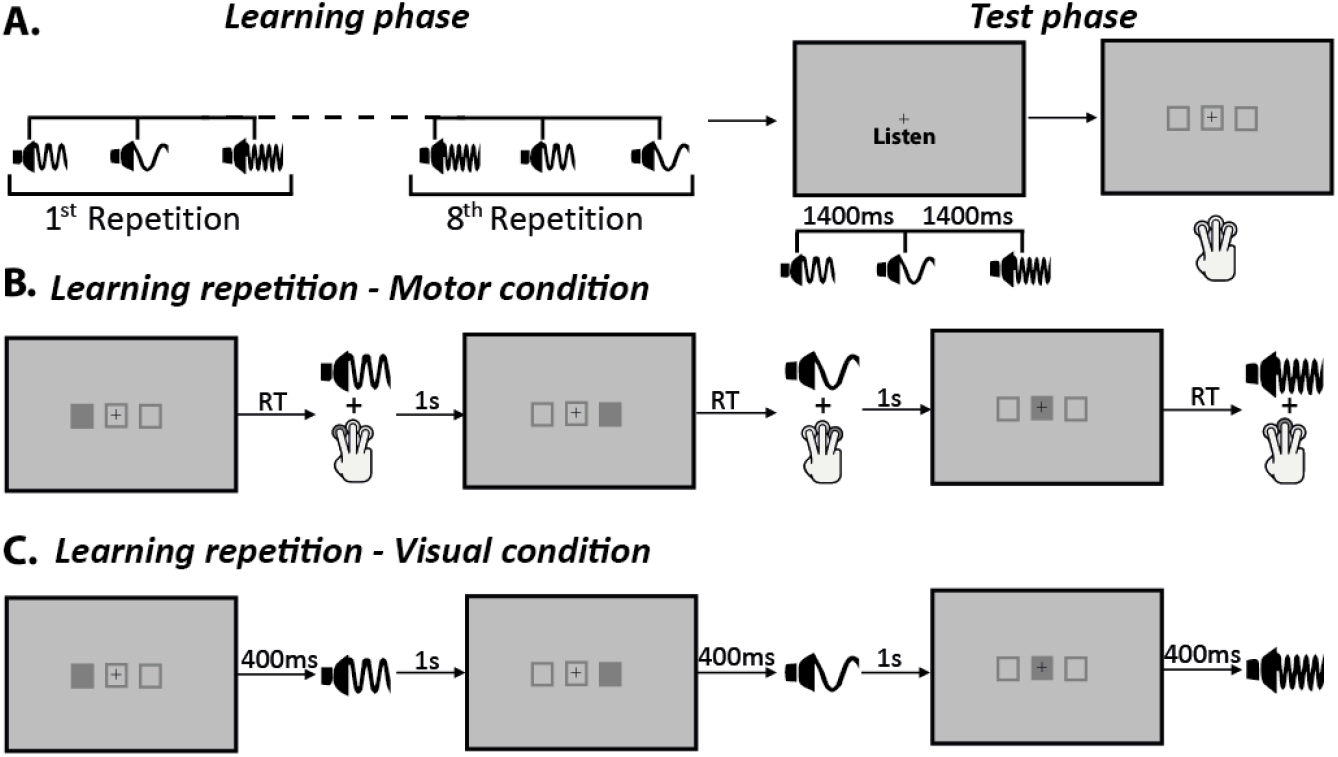
experiment design. **A** –timeline of a single experimental trial, including 8 repetitions of each cue-sound mapping in the learning phase, followed by a test phase in which participants heard a sequence of the three learned tones in a random order and had to press the buttons in corresponding order based on the button-tone mapping they acquired during the learning phase. **B** – a single learning repetition of three sounds in the Motor Cues condition. Order of cue-stimulus presentation within each repetition was randomized. **C** – a single learning repetition in the Visual Cues condition.

After the learning phase, participants were engaged in a short test (test phase) to ensure they managed to learn the correct coupling between button and tone. In this phase, participants were first presented with the three tones they had just learnt in a random order and were requested to press the associated buttons in the corresponding order, as if they were replaying the tones (see figure 1A). The correct response in the test phase was determined by successfully pressing the correct order of buttons corresponding to the three tones. Trials in which participants failed to reproduce the correct order of button presses in the test phase were discarded from analysis.

In order to examine the role of learning on the N100 sensory attenuation effect, participants also engaged in a similar learning task between visual cues and tones (Visual Cues condition). This task was also divided into learning and test phases. During the learning phase, participants were instructed to learn the coupling between a specific rectangle on the screen and the tone that followed, without pressing any buttons. Unbeknown to participants, across trials the mapping between the rectangles and the tones was identical to the mapping between buttons and tones in the Active condition. The grey rectangle always preceded the associated tone onset by 400ms, in order to simulate the reaction times in the Active condition (see figure 1C). In the test phase, participants were instructed to listen to a sequence of the 3 tones and replay it by pressing the buttons corresponding to the grey rectangles on the keyboard in the correct order (as in the motor learning condition).

Each trial in both conditions included a combination of three tones randomly drawn from the following tones: 250, 400, 600, 900 and 2000Hz. Tones were randomly assigned to each button/visual cue, with the limitation that consecutive trials will include at least one different tone from the previous trial. Each experimental block consisted of either Motor or Visual cues trials. The order of blocks was counter-balanced across participants, with half the participants starting with all the Motor blocks and the other half starting with all the Visual blocks. In total, participants completed 6 blocks of 8 de-novo learning trials from each condition (total 48 trials per condition). Before each condition, participants performed a practice of 2 trials in order to verify they understood the task. During these practice trials, participants received feedback about their performance on each trial, showing a ‘V’ if the participant responded correctly in the test phase, or an ‘X’ if responded incorrectly. During the rest of the experiment, participants did not receive performance feedback after every trial, but at the end of each block participants received feedback about the percent of correct trials throughout the previous block. This was done to encourage participants to be engaged in the task and perform the task to the best of their ability. Participants with low accuracy (less than 50% correct) were excluded completely from analysis due to insufficient data (3 participants). The experiment was programmed using Psychtoolbox-3 (www.psychtoolbox.org) on MATLAB 2019b (The MathWorks, Inc., Natick, Massachusetts, United States). Sounds were delivered using E-A-RTONE GOLD pneumatic insert earphones, with a Creative Sound Blaster Aurora AE-5 sound card.

#### EEG recording and pre-processing

EEG data were recorded using a BIOSEMI Active II 64-channel system (BioSemi, Amsterdam, the Netherlands). All electrodes were placed on the scalp using Electro-Cap according to the 10-20 system and referenced to the common model sense (CMS) electrode with an additional drive right leg (DRL) electrode serving as the ground. EEG signal was recorded at a sampling rate of 256 Hz. Four additional channels were used to record activity related to ocular artifacts (one next to each eye, one above and one below the right eye) and 2 more channels placed on right and left mastoids served as offline reference channels. EEG data were analyzed using MATLAB 2023b and EEGLAB toolbox, version 14.0.1 (Delorme and Makeig, 2004). The continuous data were re-referenced to the average of the mastoid channels and offline filtered with a 0.05-40 Hz band pass filter implemented in EEGLAB toolbox. We segmented the data into epochs covering the time window from -200ms to 1s relative to tone onset (time 0 corresponds to a button press + tone in the Motor Cues conditions and tone delivery in the Visual Cues condition). We compared the signal between conditions and repetitions using baseline correction to the average signal at -200-0ms pre sound onset (as commonly used in the literature, e.g. see (Timm et al., 2013)). For removing ocular artifacts, we used independent component analysis (ICA) as implemented in the EEGLAB toolbox. Ocular ICA components were identified by visual inspection and deleted from the global signal. Noisy trial epochs (exceeding ±100 µV range) at Fz, FCz and Cz channels were identified in the raw data and discarded from further analysis.

#### Time-frequency analysis

To further examine the influence of specific expectations on the EEG data, we used a time-frequency analysis to compare data from the first and last repetition in both the Motor and the Visual cues conditions. Time-frequency decomposition was conducted for frequencies in the 6-40Hz range. For this analysis we used a larger time window than was used for the N100 analysis, ranging from -700ms prior to tone delivery to 1s after tone delivery. All preprocessing steps were repeated for this time window. The time-frequency decomposition was calculated using the EEGLAB toolbox, with a Morlet sinusoidal wavelet transformation initially set at 4 cycles, increasing linearly to 8 cycles at 40Hz. Data were baseline corrected to the time window between -700 and -400ms prior to tone onset. The data were decomposed with a frequency resolution of 0.34Hz and 200 time points. This analysis was performed separately for data from the first and last repetition, both in the Motor Cues and Visual Cues conditions.

#### Statistical analysis

Based on previous literature, the analysis focused on the N100 component (for review see (Horvath, 2015)). N100 amplitude was defined based on the minimal value in the average signal of each participant at the time window between 50-200ms after sound onset. To examine the contribution of expectation (repetition) as well as other motor related signals (cue type) on the amplitude of the N100 component and the interaction between the two, we used a 2 (cue type: Motor / Visual) X 8 (repetition) repeated measures ANOVA. Analysis was performed only on data from trials in which participants responded correctly in the test phase. Statistical analysis of the N100 amplitude was conducted using JASP (JASP Team, 2019. Version 0.18.2.0).

Statistical analysis of the time-frequency data was conducted using a non-parametric cluster-based approach (Maris and Oostenveld, 2007). Analysis was performed on data averaged from electrodes Cz, FCz and Fz for each condition - Motor Cues 1^st^ repetition, Motor Cues 8^th^ (last) repetition, Visual Cues 1^st^ repetition and Visual Cues 8^th^ repetition. First, we performed a two-tailed paired t-test, comparing at the group level between each pair of conditions and defined time-frequency clusters showing significant (p<0.05, uncorrected) difference. For each cluster we calculated its size - the sum of t-values at each cluster. Next, we used a permutation test as described in Maris and Oostenveld (2007) to create a sampling distribution of cluster sizes and compared the real cluster sizes to this distribution. Significant clusters were defined as clusters larger than 5% of the clusters in the distribution (p<0.05).

## Results

In the current study, we examined how the auditory N100 component changes during motor-sensory (Motor Cues condition) and sensory-sensory (Visual Cues condition) de-novo contingency learning. First, we examined whether there are behavioral differences in learning levels between the two cue conditions. To this end, we examined the average performance on the tests between the Motor cues (*M*=83.8% correct, *SD*=11.9%) and Visual Cues (*M*=85.8% correct, *SD*=12.1%) conditions using Student`s t-test and Bayesian analysis. No significant difference between the two conditions was found (*t*=-1.06 *p*=0.32, *BF*_*01*_=3.21). This implies that differences in the EEG signal between the conditions cannot be explained by differences in task difficulty or performance. We also examined the reaction time (RT) of participants’ sound triggering actions during learning in the Motor cues condition. On average, participants pressed the button to trigger a sound 532.15ms after the visual prompt was presented (range 330.41-1257.48ms). RTs did not change with learning and remained similar across repetitions (1X8 Repeated Measures ANOVA, *F(3*.*46)*=0.89, *p*=0.46; result was corrected using Greenhouse-Geisser correction due to violation of sphericity).

Examining the amplitude of the auditory N100, we used a 2X8 repeated measures ANOVA with cue type (Motor Cues / Visual Cues) and repetition (1 – 8) in the learning phase as independent variables. In this analysis, a main effect of cue type is equivalent to the classic N100 attenuation effect. A main effect of repetition would be indicative of an influence of expectation learning on the N100 amplitude, and an interaction effect would suggest that the two components are dependent. We found a significant main effect of cue type, such that the amplitude of the N100 component was overall less negative in the Motor Cues condition (*M*= -6.29 *SD*= 0.52 μV) than in the Visual Cues condition (*M*= -7.89 *SD*= 0.60 μV; *F(1,29)=* 8.70, *p*= 0.006). We also found a significant main effect of repetition, such that the amplitude of the N100 was less negative as learning progressed (*F(7,203)=* 3.76, *p*= 9.37*10^-4^; see figure 2A for evoked responses in first/last repetitions and 2B for data across all repetitions). Examining the post-hoc comparisons of this analysis, we found that the main effect of learning is mostly driven by the difference between the first three repetitions and the 8^th^ repetition (1^st^ vs. 8^th^ *t*(29)= -3.34 *p*= 0.027, 2^nd^ vs. 8^th^ *t*(29)= -3.14 *p*= 0.051, 3^rd^ vs. 8^th^ *t*(29)= -3.45 *p*= 0.02, all post-hoc comparisons were corrected for multiple comparisons using Bonferroni-Holm correction). Interestingly, we did not find an interaction between cue type and repetition (*F(7,203*)= 0.55, *p*= 0.80) – suggesting that the difference between cue types remains similar during learning. Further examining this lack of difference using Bayesian analysis, we found that the model including only the two main effects had a Bayes factor of 139.87 relative to the model including the two main effects and also an interaction effect, suggesting it reflects the data better. Finally, to examine directly whether a sensory attenuation effect can be found without specific expectation, we focused on the N100 amplitude in the first repetition before any specific cue-sound association can be formed. Direct comparison of the N100 amplitude of the Motor and Visual Cues conditions in that repetition still revealed a significant sensory attenuation effect (Motor Cues: *M*= -6.55 *SD*= 3.26; Visual Cues: *M*= -8.35 *SD*= 3.30; *t*(29)= 2.93, *p*= 0.007). Taken together, these results suggest that level of cue-stimulus association affects the amplitude of the N100 component but not the sensory attenuation effect in short term learning.

**Figure 2:**
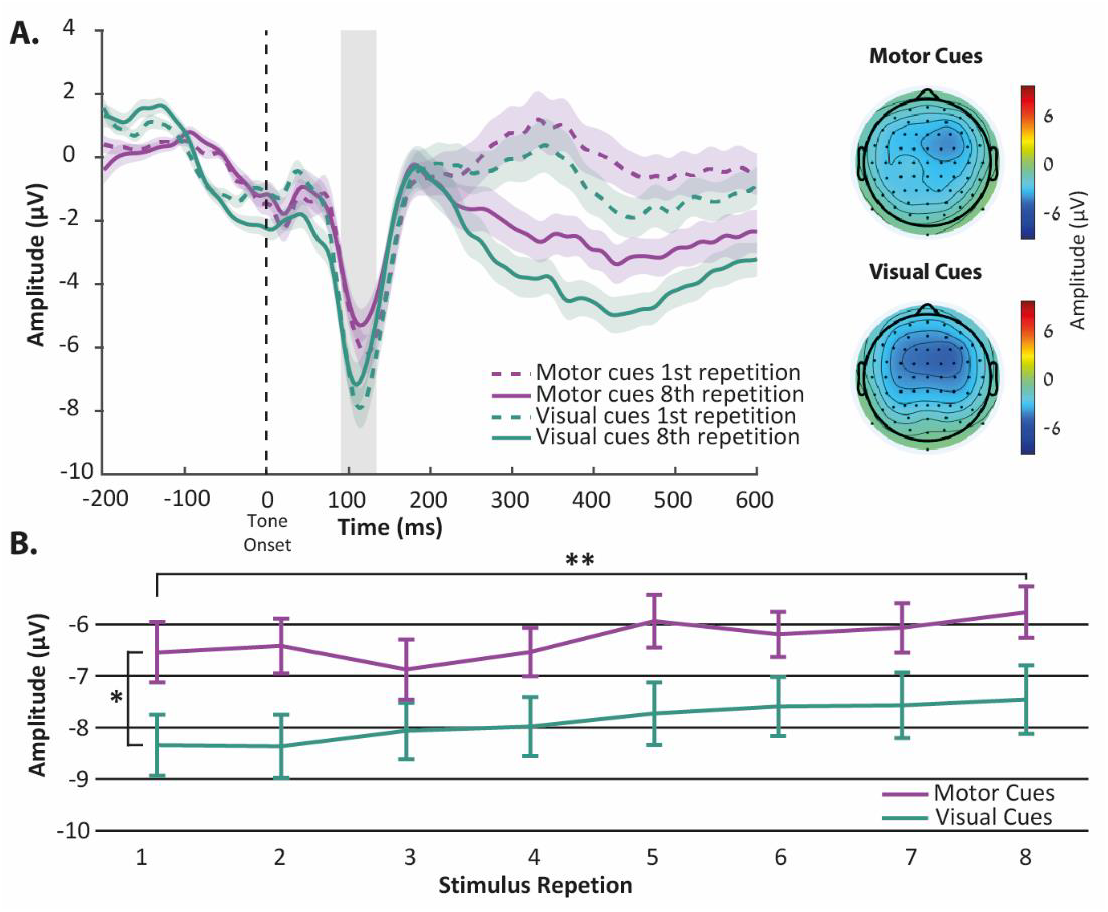
**A**. <u>Left panel</u> - Event Related Potentials (with SEM) at Cz FCz and Fz electrodes (averaged). Purple lines – Motor Cues, Teal lines – Visual Cues. Dashed lines – first learning repetition (no specific expectation), solid line – final learning repetition (after expectations are formed). Time 0 represents tone onset. Grey area represents the N100 time window. <u>Right panel</u> – topography at 100ms in all 64 channels in the Motor Cues (top) and Visual Cues (bottom) conditions. **B**. N100 amplitude in the Motor Cues (purple) and Visual Cues (teal) at each learning repetition (mean and SEM). *p<0.05, **p<0.001.

Even though the focus of our study was to examine the effects of learning on the N100 components, we next performed an exploratory analysis examining the entire time-window between 0-600ms. To establish which additional EEG components are affected by expectations, we examined differences between the first and last (8^th^) repetition, collapsed across cue types. We did so by using Student’s t-test without correcting for multiple comparisons. We found two significant time windows separating between repetitions: 113.28ms - 160.16ms (compatible with the N100 results we previously showed) and 226.56 - 600ms, compatible with an effect of repetition on the P300 and later components. To further examine the effect on the later time window (226.56-600ms), we conducted a 2X8 repeated-measures ANOVA, similar to the one conducted on the N100 data. As can be expected, we found a main effect of repetition, showing an overall more negative response as learning progresses (*F(3*.*01,87*.*21)=* 16.12, *p*= 1.97*10^-8^; Result was corrected using Greenhouse-Geisser correction due to violation of sphericity). Examining the post-hoc comparisons of this analysis, we found that the main effect of learning is mostly driven by the difference between the first repetition and 2^nd^-8^th^ repetition (1^st^ vs. 2^nd^: *t(29)*= 4.56 *p*= 1.93*10^-4^; 1^st^ vs.3^rd^: *t(29)*= 7.81 *p*= 7.69*10^-12^; 1^st^ vs. 4^th^: *t(29)*= 8.40 *p*= 2.13*10^-13^; 1^st^ vs. 5^th^: *t(29)*= 6.94 *p*= 1.18*10^-9^; 1^st^ vs. 6^th^: *t(29)*= 7.22 *p*= 2.40*10^-10^; 1^st^ vs. 7^th^: *t(29)*= 7.41 *p*= 8.52*10^-11^; 1^st^ vs. 8^th^: *t(29)*= 8.43 *p*= 1.75*10^-13^) and 2^nd^ repetition and 3^rd^, 4^th^ and 8^th^ repetition (2^nd^ vs.3^rd^: *t(29)*= 3.25 *p*= 0.026; 2^nd^ vs. 4^th^: *t(29)*= 3.83 *p*= 0.003; 2^nd^ vs. 8^th^: *t(29)*= 3.87 *p*= 0.003; all post-hoc comparisons were corrected for multiple comparisons using Bonferroni-Holm correction). We also found a main effect of cue type, such that the Motor cues condition elicited a less positive response relative to the Visual cues condition (*F(1,29)*= 5.94 *p*= 0.021) and no interaction effect between Cue type and repetition (*F(7,203)*= 0.50 *p*= 0.83). These results are in agreement with the N100 results, suggesting that both Repetition and Cue type affect the P300 and later components independently.

Finally, we examined the differences between cue types and repetitions in the time-frequency domain. To this end, we used a cluster-based approach to define clusters differentiating between the two conditions (see methods). First, we compared the time-frequency maps between the first and last repetition in the Motor and Visual Cues separately, to evaluate the times and frequencies affected by learning. In both cue conditions, we found one significant cluster distinguishing between repetitions at the time window between -400 and -200ms. At this time window, we found lower power for 12-40Hz frequencies in the last repetition compared to the first repetition (see lower panel of figure 3). Next, we examined the effect of cue type within each repetition. In both the first and the last repetition we found a significant cluster distinguishing between the cue types in the time window between -200 to 400ms, with higher power in the Visual Cues condition relative to the Motor Cues condition (20-40Hz in first repetition and 15-35Hz in the last repetition; see right panel in figure 3). This pattern of results shows that the difference between cue types (Motor / Visual) is represented on a later time-window than the difference between repetitions and further supports that these two factors affect neural activity independently.

**Figure 3:**
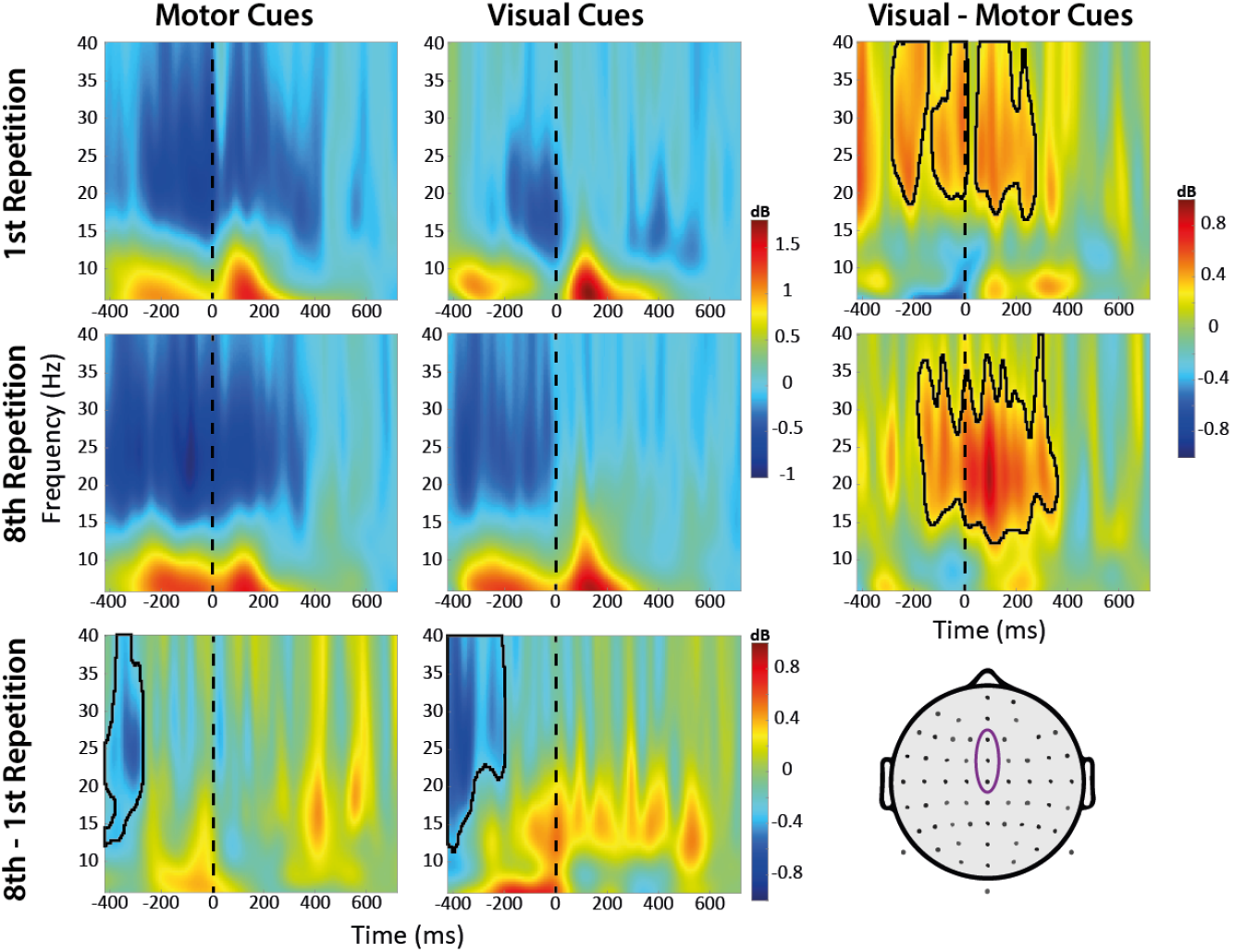
Time-frequency data for Motor and Visual Cues (columns) in the first and last (8th) repetition (rows). Lower and Right panels represent the difference between repetitions and cue types respectively. Marked areas represent significant clusters in each comparison. Dashed lines represent time of tone onset. Data was taken from electrodes Fz, FCz & Cz (bottom right panel).

## Discussion

The sensory attenuation effect is commonly explained in the literature either as driven by specific expectations of sensory outcome (Wolpert and Flanagan, 2001) or as a global effect of action-outcome binding (Weiss and Schutz-Bosbach, 2012). In the current study, we teased apart the contributions of each of these factors by examining the magnitude of sensory attenuations during short-term cue-outcome learning process. Specifically, participants learned to couple between button presses and tones and between visual cues and tones, such that at the first repetition of the coupling process participants had no knowledge about the identity of the upcoming tone. We also ensured that by the end of the learning process, participants successfully learned the coupling between cues and tones. We found that, regardless of cueing modality (Motor / Visual), N100 evoked responses decreased in amplitude as learning progressed, meaning that expectation of a specific outcome gained by experience affects the magnitude of the neural responses they evoke. We also replicated the classic sensory attenuation effect, such that regardless of learning phase, tones that were cued by an action evoked smaller N100 responses than identical tones cued by a visual event. Interestingly, we found no interaction between the two factors in affecting the magnitude of the N100 evoked responses, and specifically a significant sensory attenuation effect already at the first repetition of the learning process. This implies that sensory attenuations do not depend on prior knowledge regarding the identity of the sensory outcome. Our results suggest that while both specific expectations of sensory outcome and the type of preceding cue (Motor / Visual) affect the magnitude of sensory evoked responses, the sensory attenuation effect (difference between responses following motor and sensory cues) remains constant regardless of expectation level, suggesting it is not influenced by it, at least following short-term learning. A similar pattern of results was also observed in later evoked-response components and in the time-frequency domain, where we found that repetition and cue type influenced neural processing in distinct time-windows and frequency bands.

A novel aspect of the current study was the fact that our design allowed us to compare auditory responses following motor and visual cues during the learning process, as expectations are formed. Previous studies have shown that expectations shape sensory evoked responses (Schroger et al., 2015). This has been shown not only in the N100 component but also in the P300 component (Polich, 2007), similar to our current results. Similarly, in the motor domain, expectations have been shown to affect the evoked responses for the actions’ sensory consequences (Schroger et al., 2015; Elijah et al., 2016; Fuehrer et al., 2022; Storch and Zimmermann, 2022; Thomas et al., 2022). Since the sensory attenuation effect is the direct comparison between responses to sensory stimuli cued by motor and non-motor events, it can be influenced by expectation-dependent changes in signal amplitude in the motor-sensory condition, sensory-sensory condition or both. Previous studies examining the role of expectation on the sensory attenuation effect have only measured expectation-dependent changes in the motor condition, but not in the passive condition (Elijah et al., 2016; Fuehrer et al., 2022). However, as our current results show, evoked responses in passive conditions are also sensitive to expectation levels that need to be taken into account. Indeed, when this is the case, we find no changes in sensory attenuation as a function of expectation at least for short-term learning. To the best of our knowledge, this is the first study to take both effects into account.

Our results support the idea that the sensory attenuation effect does not depend on foreknowledge about the identity of upcoming sensory events. This suggests that the sensory attenuation effect is a global feature of sensory-motor processing that happens irrespective of experience-dependent knowledge regarding the sensory consequences of the action. This is in agreement with previous studies showing that the sensory attenuation effect occurs even when controlling for the prediction level of upcoming sensory stimuli (Weiss and Schutz-Bosbach, 2012; Dogge et al., 2019; Klaffehn et al., 2019; Storch and Zimmermann, 2022). It is important to note that in the current study, in the first repetition of learning, participants had no expectation about the identity of upcoming sounds but did have a general expectation to hear a sound at a given time. It is still unclear how sensory evoked responses are shaped in the absence of any specific expectation about modality or timing of action-stimulus coupling.

While the current study shows that the N100 attenuation effect does not depend on knowledge about the identity of upcoming events, it is still an open question what information is encoded in such signals. One prominent suggestion is that the sensory attenuation effect is related to brain mechanisms marking the presented stimulus as self-triggered. A previous study has shown that reported sense of agency (SoA) over an upcoming stimulus is mostly related to later components such as P200 and P300 (Seidel et al., 2021). Nevertheless, the N100 may still represent an overall ownership over the sensory outcome, independently of the specific level of reported SoA. Another possible type of information encoded in the N100 attenuation effect represents properties of the stimulus-triggering action. Previous studies have shown that both perception of sensory events and activity in sensory regions are modulated by the identity of the stimulus-triggering hand (Reznik et al., 2014; Buaron et al., 2020), however it is still unclear which EEG components are sensitive to such factors.

There is a debate in the literature about the underlying mechanisms of motor (internal) and sensory (external) predictions. Wolpert and Flanagan (2001) suggest that motor predictions are a unique mechanism that enables us to move and act in the world. Other predictive theories suggest that motor and sensory prediction share a common mechanism (Friston, 2010; Stekelenburg and Vroomen, 2015). The predictive coding theory (Friston, 2010) suggests that probabilistic models in the brain send their predictions to lower levels of the hierarchy and receive in return information about prediction error used to update the models. This theory suggests that many global theories of brain function, including the motor forward model (Wolpert and Miall, 1996), can be united under a unified perceptive model of the brain as a generative model of the world. Neural models suggest that predictions modulate the responsiveness and tuning of stimulus-selective neurons, such that their activity is more selective towards the expected event (Summerfield and de Lange, 2014; Yon et al., 2018). While our current results cannot resolve these debates, we do find a motor-unique component that affects the sensory evoked response. Furthermore, the lack of interaction between expectations and cue type that we report in the current study is in better agreement with the notion that expectation effects are common to motor and non-motor prediction modalities.

## Acknowledgements

The study was supported by the Israel Science Foundation (grant No. 2392/19 to R.M.). The authors thank Dema Hreaky for her assistance in data collection and all lab members for constructive comments and fruitful suggestions.

## Notes

### Competing Interest Statement

The authors have declared no competing interest.

